# Inverse Reinforcement Learning to Study Motivation in Mouse Behavioral Paradigms

**DOI:** 10.1101/2024.06.13.598948

**Authors:** Andre Telfer, Afsoon Alidadi Shamsabadi, George Savin, Junfeng Wen, Alfonso Abizaid

## Abstract

Motivation describes the underlying goals that drive animal and agent behavior. In Neuroscience, behavioral paradigms are used to quantify the motivations of mice and used to gain insights into traits and diseases which can be translated to humans. In recent years, Computer Vision models are becoming widely adopted by Neuroscientists to score mouse behavior associated with motivations such as hunger and anxiety. However, a single motivation can be expressed by multiple different behaviors, and a single behavior can be linked to multiple motivations. Therefore the ideal analysis of motivational paradigms would attempt to directly recover the underlying motivations guiding behavior, rather than indirectly score their associated behaviors. In this paper, we move towards this goal by applying Inverse Reinforcement Learning to study the underlying motivations that drive mouse behavior.

## 1. Introduction

Motivation guides behavior and is associated with many diseases such as depression and obesity, which impact a large percentage of the population. Mouse models are a common way to study human traits and diseases, and are widely employed in preclinical trials of drugs that affect motivation such as antidepressants, anxiolytics, and obesity medication. Mouse behavioral paradigms are commonly used to quantify behaviors associated with motivation and emotionality, and to compare differences in motivation to identify treatment effects. Figure 1 shows a common mouse paradigm, the Open Field Test [3], which can be used to study motivated behaviors such as those associated with anxiety and hunger. These paradigms were traditionally scored manually on simple measures relating to position and movement. For example, in Figure 1 feeding motivation can be quantified as time spent near the food, while anxiety behavior is quantified as time spent in the sides and corners of the box where, in natural environments, mice are less vulnerable to predators.

**Figure 1.**
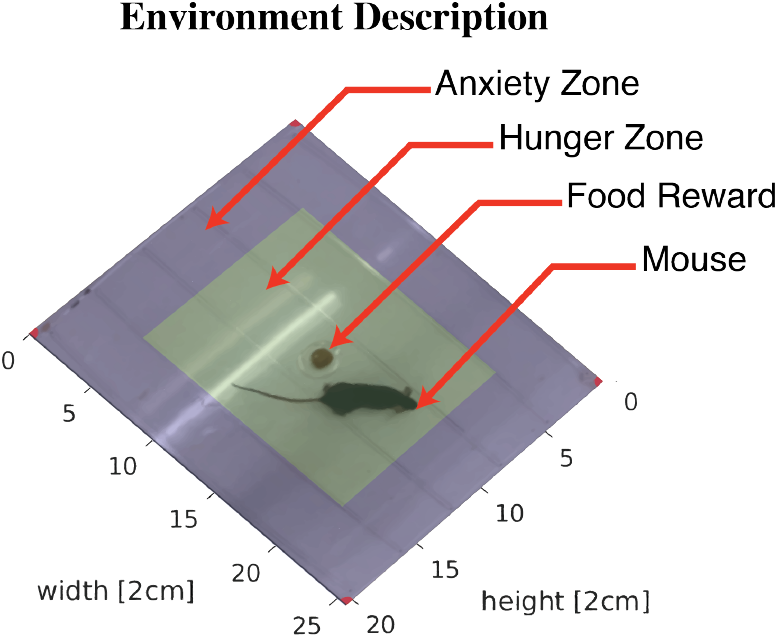
The Open Field Test is a mouse behavioral paradigm commonly used in Neuroscience where a mouse is placed in an open box and recorded to make inferences about motivation.

In this work, we use various vision tools to extract visual representations of the mice and apply Inverse Reinforcement Learning (IRL) to learn plausible rewards that can potentially explain the behavior and underlying motivation of the mice. To test this we use data from an experiment where mice have been treated with drugs that affect motivation.

## 2. Related Work

### Supervised Behavior Analysis

In recent years, there has been a swift growth in the number of AI tools available to researchers in behavioral neuroscience. These tools have been powered by advances in deep learning and have enabled accurate and fast scoring of large-scale data sets. When parts of the behavior are well defined, supervised approaches such as DeepLabcut [7] and SiMBA [9] can be employed to quantify large amounts of data. However, in many cases, we do not know all of the behaviors which are useful to score when studying motivation.

### Unsupervised Behavior Analysis

A more flexible approach to identifying behavioral patterns and differences is to employ unsupervised methods such as B-SOiD [5]. While unsupervised approaches are useful for identifying new clusters of behavior and differences between groups, it does not attempt to directly tie these behaviors to motivation. Behavioral differences can occur for a multitude of reasons outside of the focus of the study, and behaviors linked to the same underlying motivation can be grouped into separate clusters.

### Inverse Reinforcement Learning

Another approach that has not been applied to behavioral paradigms studying motivation is Inverse Reinforcement Learning (IRL). The purpose of IRL is to take observations of an agent’s behavior and identify the underlying reward the agent receives that would explain the actions they took. While current methods of studying motivation are indirect through quantification of behavior, IRL directly aims to identify the reward of agents that explains their motivations and behavior. An example where we expect IRL rewards to be different than behavior is with movement towards a motivated goal. A simple behavior analysis tracking the position of the animal will put equal weight on each location it passes through, however, IRL will emphasize the end goal that better reflects their motivation.

## 3. Methods

### 3.1 Mouse Experimental Data

The mouse behavioral data was taken from an existing study that examined the anxiolytic effects of Ghrelin (a hormone associated with increased feeding motivation [1]), and its interaction with Melanotan II (MTII) which passively opposes the effects of Ghrelin on hunger [6]. We will refer to the Ghrelin group as being *hunger* motivated, while the MTII group as being *anxiety* motivated (indirectly caused by a decrease in the hunger motivation). The study consisted of four groups of 20 mice that were given the following treatments:

- Control: two saline injections.
- Ghrelin: one saline and one Ghrelin injection.
- MTII: one saline and one MTII injection.
- MTII-Ghrelin: one MTII and one Ghrelin injection.

After injection, the mice were then placed in a 40×50cm Open Field box with a piece of food (3g of cookie dough) placed in the center (Figure 1) and were recorded for 10 minutes at 30 fps. The time spent near the food is taken as a behavioral indicator of hunger motivation while spending time near the edges/corners of the box is considered an anxiety-like behavior. It’s important to note that the control group does not exclude a natural hunger motivation. In this experiment, we observe that the Control group can exhibit behavior similar to the Ghrelin group.

### 3.2 Preparing Gridworld Environment for IRL

To track the mouse in the environment we trained a pose estimation model to identify body parts (nose, ears, neck, midbody, and tail). To create a dataset, we extracted 200 frames from a sample of 5 videos, manually labeled the position of body parts in frames, and then trained a pose estimation model using DeepLabCut [7] with a ResNet50-based architecture [4].

To combine data from videos that had different camera perspectives and box positions, we transformed the tracking data to a common frame of reference (see Figure 2). The homogeneous transformation to move between these two coordinate systems was computed by manually labeling the box corners in each video and matching them to corresponding points on a box schema (40×50cm). This transformation could then be applied to all of the mouse body part coordinates previously extracted. Next, we binned the mouse centroid (mean position of all body parts) into a 20 × 25 grid of 2cm squares. The state of the environment was represented as the 20 × 25 matrix of 0s, with 1 representing the mouse centroid. (*n* = 500 possible position states). To identify the agent’s actions, we looped over the states in order and determined whether the agent stayed still or moved horizontally or vertically (*k* = 5 possible actions). In the rare occurrence that the mouse/agent jumped between non-adjacent states, an artificial sequence of actions was constructed between the start and the end states. To simplify later steps, the state consisted only of one single *x, y* position (midbody). The result was a dataset of 1.4 million agent state-action pairs extracted from 80 mice over 12h of video. The final environment was implemented as a Gymnasium environment [2].

**Figure 2.**
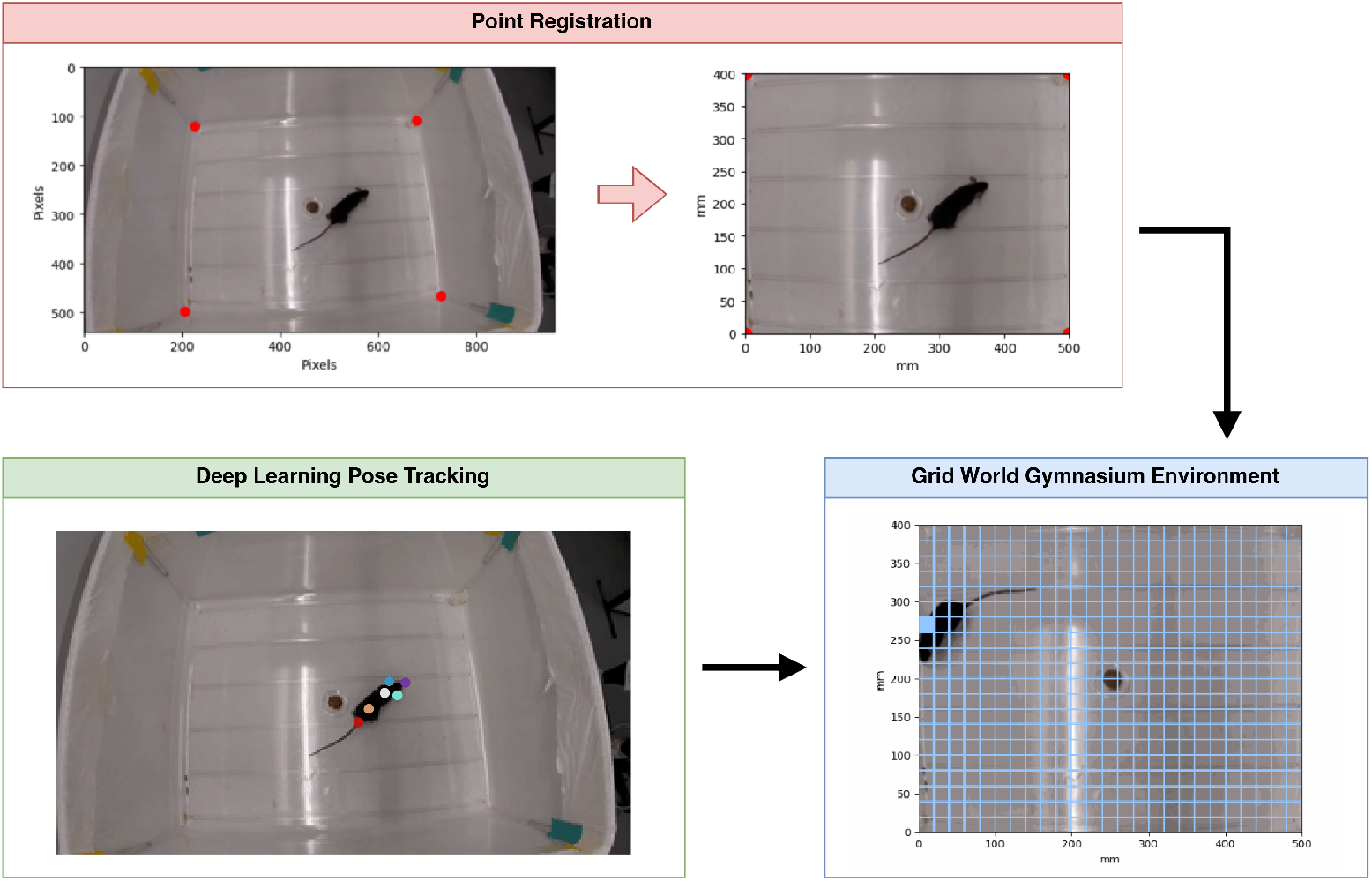
We created a simple environment for performing IRL by tracking the position of the mouse and transforming it to fit in a grid world with 20x25 squares.

### 3.3 RL with Linear Programming

For the IRL method we use Linear-Programming Inverse Reinforcement Learning (LP-IRL) [8]. LP-IRL uses the sampled trajectories of a Markov Decision Process (MDP) and solves the reward function as a linear combination of the state features. In other words, the reward of state *s*, can be represented as *R*(*s*) = ***α***^*T*^ ***ϕ***(*s*) where ***ϕ***(*s*) *∈ ℝ*_*d*_ represents the feature vector for state *s*, and ***α****∈ ℝ*_*d*_ represents the variables which are supposed to be found by the algorithm. Defining 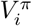 as what the state value function of the policy *π* in the MDP would have been if the reward function had been *ϕ*_*i*_, the actual state value function of the policy *π* will then be the linear combination

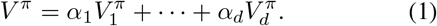

In this regard according to [8], the variables ***α*** can be obtained by solving the following optimization problem

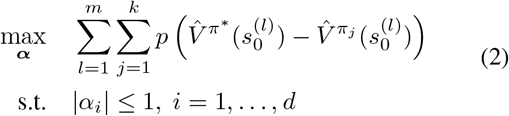

where *p*(*x*) = *x*, if *x* ≥ 0, and *p*(*x*) = 2*x*, if *x* < 0, π^*^ is the assumed optimal policy from which the agent’s sampled trajectories are available, π_*j*_, ∀*j* ∈ {1, …, *k*} represent the *k* random policies and 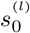 is the starting state of the *l*^th^ trajectory (*l* = 1, …, *m*). Intuitively, problem (2) encourages the agent’s behavior policy π^*^ to be better than random policies π_*j*_ for the underlying MDP. It can be cast as a linear program and the algorithm is given in Algorithm 1.

#### Algorithm 1

Algorithm of LP-IRL implementation.

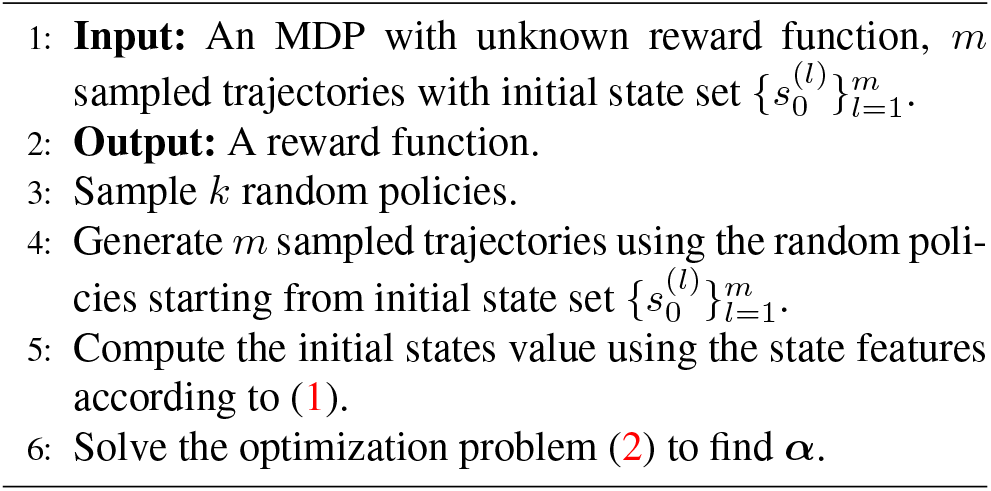

To apply the LP-IRL method to our setting, the state trajectories of the mice within the same group were segmented into intervals of 300 non-overlapping frames. The LP-IRL method was then applied to these trajectories to extract rewards. Finally, the learned rewards were then normalized to [0, 1]. Results are reported as the average over 10 runs. Experiments were conducted with γ = 0.9.

## 4. Results and Analysis

Results of the IRL rewards are shown in Figure 3 which compares the behavior of mice in each treatment group, and the IRL-extracted rewards which offer plausible explanations of their behavior. The top row shows the average state visitation frequencies of mice, and the bottom row shows the extracted rewards in each group.

From the top row, one can observe that all four groups exhibit a strong inclination of staying at corners, which is considered an anxiety-like behavior under the design of the Open Field Test [3]. From the biological perspective, the largest expected differences between groups are between the MTII vs Ghrelin groups, which have been given treatments that impact motivation in opposing directions (oppressing vs incentivizing hunger). These expected differences were reflected in both the behavior and reward plots in Figure 3. The visitation and reward of the MTII group around the center where the food was placed are significantly lower than those in the Ghrelin group. The MTII group has noticeably higher values in the corners than the Ghrelin group, showing a noticeable anxiety effect from the behavior and RL perspectives. In addition, the reward plots are generally more distributed compared to the behavioral plots, which could be explained as some exploration bonus for the mice to familiarize the new environment.

**Figure 3.**
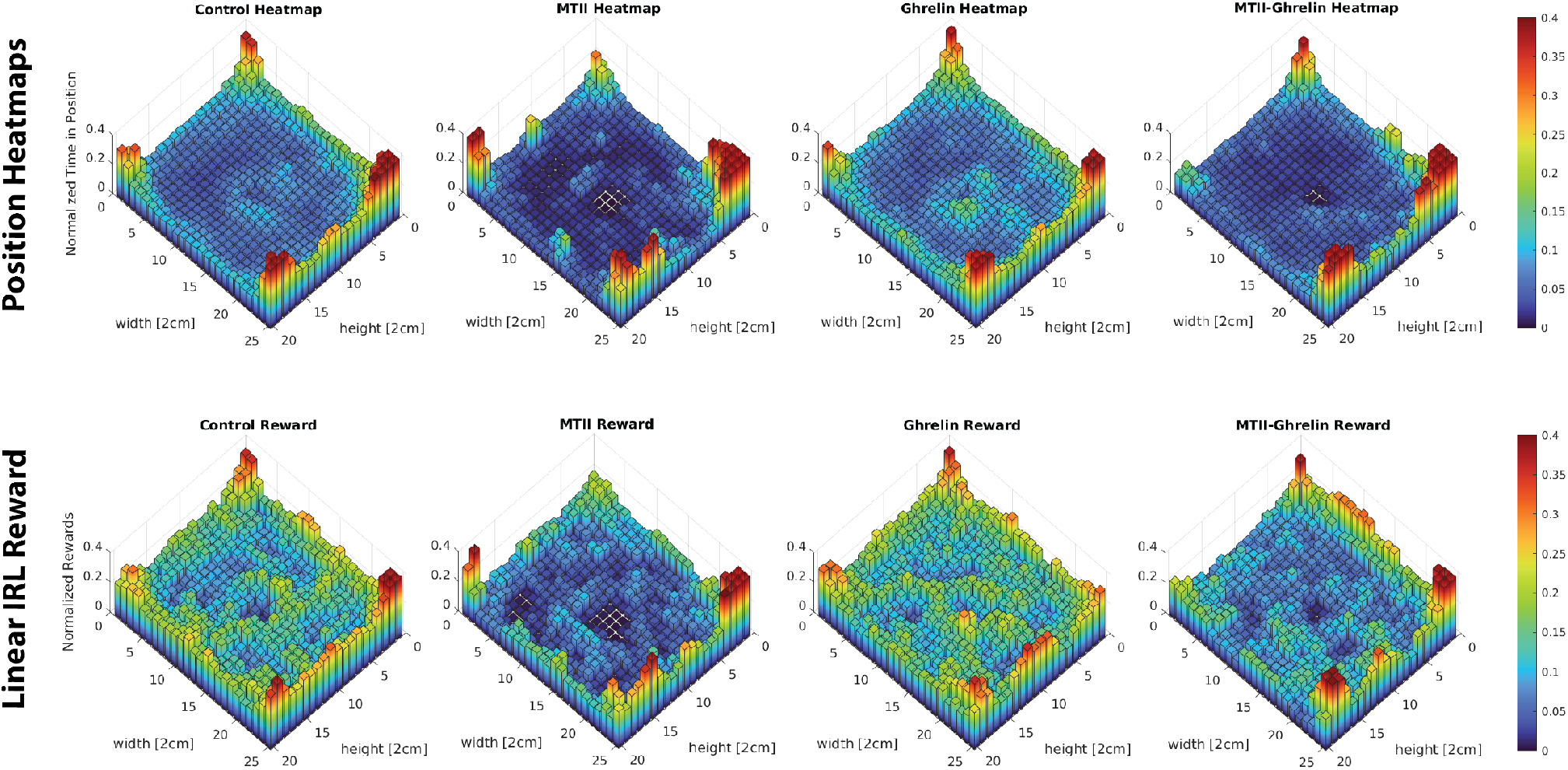
The top row represents the state visitation frequencies (where the mice spent their time), while the bottom row displays IRL-extracted rewards a mouse received for being in a certain position. Since all groups displayed a strong corner preference, we visualized the differences in hunger motivation by setting the z-axis and color bar upper-limit to 0.4.

The primary difference between the behavior (Figure 3 top row) and the reward (Figure 3 bottom row) is that the normalized reward in the center was higher compared to the time spent in the center (Figure 1), furthermore, we see a more uniform baseline reward across the Open Field box. These elevated center rewards indicate that mouse behavior is partially described by short-term reward-seeking behavior, rather than just the long-term return of the more optimal corner states. The uniform baseline may also be seen as evidence of an exploration motivation.

## 5. Further Discussion

Our observations assume that the position-based state representation, which is currently scored in traditional behavioral analysis, provides enough information to model rewards. In the future, solving for rewards at separate time intervals or utilizing a more complex state representation could be explored. Binning over time intervals could better explain observed characteristics of mouse behavior such as acclimation, which describes diminishing anxiety over prolonged exposure to a novel environment. Similarly, keeping information about previously visited states may be helpful when modeling exploration motivation.

The research direction of modeling agent rewards opens up new possibilities to validate our understanding of motivation through RL agents. In Cognitive sciences, models of the mind are sometimes validated by creating agents that use them to solve problems. This IRL approach for extracting information about agent motivation opens up a similar opportunity for behavioral neuroscience, as RL agents can be trained to maximize the reward they receive from an environment similar to real mice. How closely these simulated behaviors match the real behaviors is a reflection of how well we have modeled the agent motivation.

## 6. Conclusion

In Neuroscience, mouse models are commonly used to study human traits and diseases, including those related to motivation. These paradigms traditionally score behaviors associated with motivation. However, the mapping between motivation and behavior is not 1-to-1 and a more ideal approach would be to directly infer mouse motivation. In this work, we apply Inverse Reinforcement Learning to a mouse behavioral paradigm in order to infer plausible motivations guiding mouse behavior. These rewards capture contrasts between treatment groups have different properties than those achieved by standard behavioral analysis.

This IRL approach to modeling behavior is not intended to directly replace statistical methods of behavior analysis. However, it opens up future possibilities for validating our understanding of motivation and the interaction between interventions by training and observing the behavior of RL agents based on the IRL rewards.

